# Modeling bottlenecks, modularity, and context-dependency in behavioral control

**DOI:** 10.1101/2021.08.06.455452

**Authors:** Anjalika Nande, Veronika Dubinkina, Riccardo Ravasio, Grace H. Zhang, Gordon J. Berman

**Affiliations:** Department of Physics, Harvard University, Cambridge, MA 02138, USA; Institute for Computational Medicine, Johns Hopkins University, Baltimore, MD 21218, USA; Department of Bioengineering and Carl R. Woese Institute for Genomic Biology, University of Illinois at Urbana-Champaign, Urbana, IL 61801, USA; Institute of Physics, École Polytechnique Fédérale de Lausanne, 1015 Lausanne, Switzerland; James Franck Institute, University of Chicago, Chicago, IL 60637, USA; Departments of Biology and Physics, Emory University, Atlanta, GA 30322, USA

## Abstract

In almost all animals, the transfer of information from the brain to the motor circuitry is facilitated by a relatively small number of neurons, leading to a constraint on the amount of information that can be transmitted. Our knowledge of how animals encode information through this pathway, and the consequences of this encoding, however, is limited. In this study, we use a simple feed-forward neural network to investigate the consequences of having such a bottleneck and identify aspects of the network architecture that enable robust information transfer. We are able to explain some recently observed properties of descending neurons – that they exhibit a modular pattern of connectivity and that their excitation leads to consistent alterations in behavior that are often dependent upon the prior behavioral state (context-dependency). Our model predicts that in the presence of an information bottleneck, such a modular structure is needed to increase the efficiency of the network and to make it more robust to perturbations. However, it does so at the cost of an increase in context-dependency. Despite its simplicity, our model is able to provide intuition for the trade-offs faced by the nervous system in the presence of an information processing constraint and makes predictions for future experiments.

---

When presented with dynamical external stimuli, an animal selects a behavior to perform — or a lack thereof — according to its internal drives and its model of the world. Its survival depends on its ability to quickly and accurately select an appropriate action, as well as to transmit information from the brain to its motor circuitry in order to physically perform the behavior. In almost all animals, however, there exists a bottleneck between the number of neurons in the brain that make cognitive decisions and the motor units that are responsible for actuating movements, thus constraining the amount of information that can be transmitted from the brain to the body^1,2^.

In the fruit fly *Drosophila melanogaster*, descending commands from the brain to the ventral nerve cord (VNC) are transmitted through approximately 300 bilaterally symmetric pairs of neurons that have their cell bodies in the brain and have axons project into the VNC^3,4^. Recent anatomical studies have shown that these neurons exhibit a modular pattern of connectivity, with the descending neurons clustering into groups that each innervate different parts of the motor system^5,6^.

In addition to these anatomical properties, in the fruit fly, manipulating these descending neurons via optogenetics has shown that exciting individual neurons or subsets of neurons often result in dramatic and robust behavioral alterations – for example, exciting the DNg07 and DNg08 neurons reliably elicits head grooming, and exciting DNg25 elicits a fast running response^7^. In many cases, however, it has been shown that exciting the same neuron in different contexts (e.g., walking and flying) often have context-dependent effects^7–9^. In other words, the behavioral effect of stimulating the neuron often depends on the actions that the fly was exhibiting just prior to the activation.

In this study, we use a simplified model of behavioral control to explore how modularity may help increase the efficiency and robustness of behavioral control given an information bottleneck. Specifically, our model predicts that modularity increases the efficiency of the network and its robustness to perturbations, but also that modularity increases the amount of context-dependency in how behavioral commands are transmitted through the bottleneck. While our feed-forward model is a vast over-simplification of the complicated recurrent circuitry that lives within a fly’s ventral nerve cord, we show that it provides intuition into the trade-offs the nervous system has to navigate and makes qualitative predictions as to how the system might respond to inhibition or double-activation experiments.

## RESULTS

Inspired by the fly ventral nerve cord, we have developed an abstracted model that aims to generate insight into the general problem of behavior control through an information bottleneck. Specifically, we assume that there is a set of *N* behaviors that are in an animal’s behavioral repertoire and that to perform one of these behaviors, the animal must excite a subset of *M* total binary “motor” neurons (e.g., task 14 requires units 1, 3, and 99 to turn-on, and all the rest to be turned off - see Fig. 1a-b). However, to model the effect of having limited information transmission from the brain to the motor systems, any commands from the brain must travel through an intermediate layer of *R* < *M, N* descending neurons^5^. We implemented this model using a feed-forward neural network, with the task being encoded in the top layer, the descending neurons being the intermediate layer, and the motor units constituting the bottom layer (see Fig. 1a). For simplicity, we assume that the brain’s intended behavioral output is represented in a one-hot encoded manner, where only one “decision” neuron is turned on at once (i.e., behavior 2 is represented by a first layer of {0, 1, 0, …} ∈ {0, 1} ^*N*^). We start with the case where each behavior is randomly assigned a set of *k* motor neurons that must be activated. Fig. 1b shows an example of this desired mapping, which we call our behavioral matrix. To perform a behavior, one of the decision neurons has to be activated and pass its signal through the network. The parameters of the network, weights 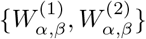 and biases 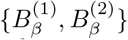, are trained to perform the mapping between the top and bottom layers as accurately as possible (see details in Methods).

**FIG. 1.**
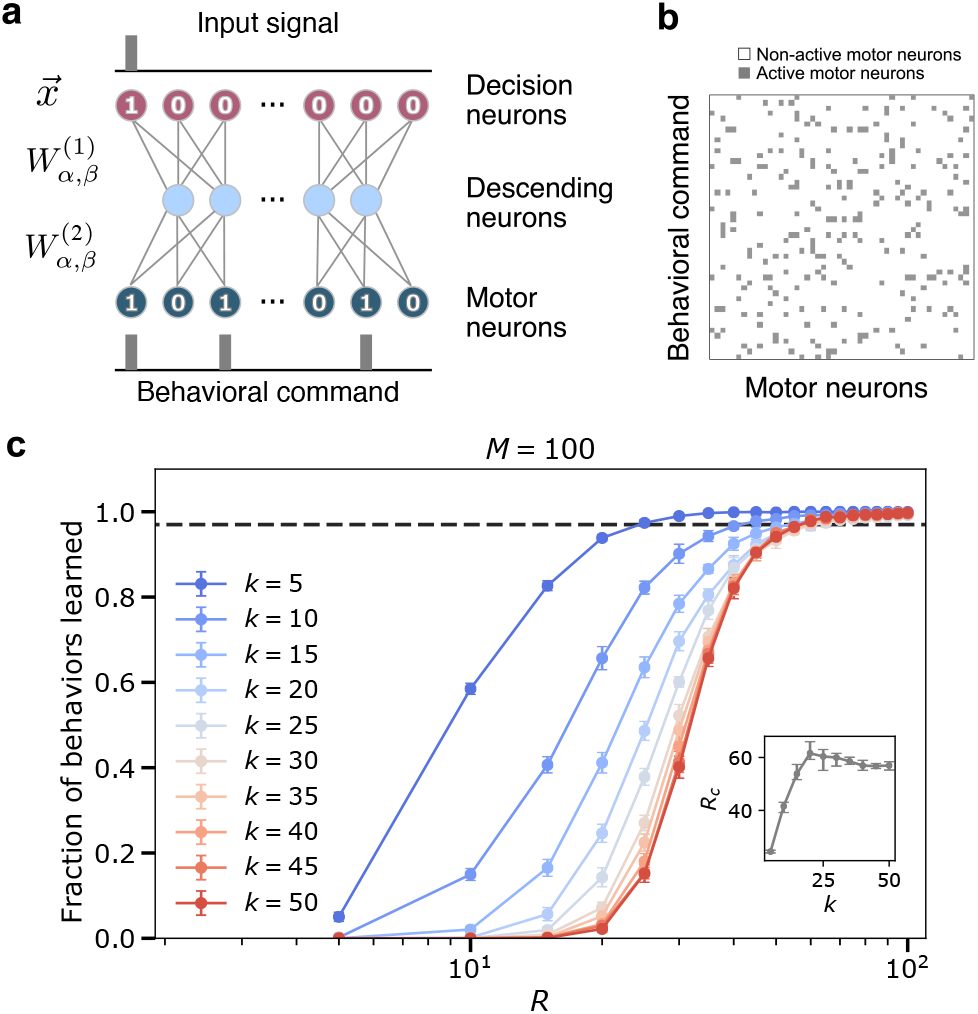
Model construction and parameters. **(a)** The structure of the ventral nerve chord is modeled by a neural network that takes as input a task assignment represented by a binary sequence 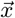 of length *N*. The signal travels through a hidden layer (size *R*) to an output layer (size *M*), which corresponds to descending neurons and motor neurons respectively. Each neuron in one layer communicates with all the neurons in the following layer through the weight matrices 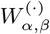, tailed in Methods. **(b)** An example of a behavioral matrix that indicates the motor units activated for each task. Row *i* corresponds to the *i*-th behavioral command (i.e., the *i*-th neuron activated in the input layer of the network). *k* is the number of motor neurons needed to execute a given behavior. Columns correspond to different motor neurons (i.e., the *j^th^* column indicates whether a particular motor neuron was active (grey) or not (white) in the behaviors). **(c)** Fraction of behaviors learned as a function of hidden layer size *R* and fixed input layer size *N* = 100 for varying *k* and fixed out-put layer size *M* = 100. The inset shows the critical bottle-neck size *R_c_* as a function of *M*. Each point is averaged over 30 random input-output combinations. Dashed line indicates critical bottleneck threshold.

Given this model, we would like to study how the network performs as a function of the bottleneck size and the sparsity of the behavioral matrix. The absolute maximum number of sequences that the network could encode is 2^*R*^. However, this simple neural network is incapable of reaching the ideal limit. In Fig. 1c, the bottleneck size required for accurate encoding is 20 60 for *N* = *M* = 100, depending on the sparsity of the behavioral matrix. These values are much larger than the minimal possible bottleneck size, *R* = log_2_ 100 ≈ 7. While we will explore the potential reasons for this discrepancy shortly, we empirically define the critical bottleneck size, *R_c_*, as the minimal number of neurons in the intermediate layer sufficient to reproduce 98% of the behaviors correctly, averaged across multiple random instantiations of the behavioral matrix. See Suppl. Fig S1 for example learning and loss curves.

To explore how the statistics of the behavioral matrix affects the critical bottleneck size, we altered the sparsity of the outputs by manipulating the number of motor neurons activated per behavior (*k*) while keeping *M* = *N* = 100 (Fig. 1c). Note that since our output size is 100 and its encoding is binary, a neural network with *k* and 100 − *k* activated motor neurons have the same statistical behavior. Thus, sparsity increases as *k* deviates from 50 in either direction. As evident from Fig. 1c and the inset therein, as *k* decreases below 25, the network requires fewer neurons in the intermediate layer (a lower *R_c_*) to learn all of the behaviors perfectly, with the decrease starting around *k* = 25. Ultimately, for the sparsest output encoding we tested (*k* = 5), the network requires half the number of neurons compared to the densest (*k* = 50) case (*R_c_* ≈ 24.4 ± 0.8 vs. *R_c_* ≈57 ± 2). This is because it is more difficult for our model to learn the more complicated patterns that are associated with a denser output. Furthermore, we note that the shape of the curve, as a function of hidden layer size, *R*, approaches that of a logistic function in the limit of dense output signal (as *k* approaches 50).

Equivalently, sparsity can be varied by fixing *k* and varying the size of the output layer *M* (here, keeping *N* = 100 fixed) (Fig. S2). We again find that as the output signal becomes more sparse, that is, as M increases, it is easier to learn the mapping from behavior to motor commands. Moreover, we also notice that the learning curves split into two regimes (Fig. S2a-b) corresponding to when *M* is smaller or larger than *N*. When *M > N*, the network finds it much easier to learn with the learning ability saturating when the bottleneck size is a certain fraction of the output layer.

### Modularity of behaviors

While the analyses presented in the previous section involved random mappings between behaviors and motor outputs, we now ask if imposing biologically inspired constraints on this mapping might affect the efficiency of the network. Specifically, we will assume that the behavioral matrix is modular, with similar behaviors (e.g., different locomotion gaits or different types of anterior grooming motions) more likely to require similar motor output patterns. This constraint is motivated from previous anatomical studies in *Drosophila*^5^.

To explore the effect of modular structure on our model, we performed a set of simulations with various degrees of behavioral matrix modularity. Specifically, we fixed *k* = 10 and split the behavioral matrix into 5 regions (see inset in Fig. 2a). If there is no active motor neuron in common between the different clusters, then we have perfect modularity (*μ* = 0.8, where *μ* is the fraction of the edges that fall within the modules minus the expected fraction within the modules for an equivalent random network^11^, see Methods). We then allowed for some overlap between regions to generate matrices with a spectrum of modularities between the perfect modular limit and random mixing. We observed that the modular behavioral matrices can be learned more efficiently than random matrices, requiring far smaller critical bottleneck sizes to achieve the correct mapping of behavioral commands (Fig. 2a). The perfectly modular output matrix was learned with only *R_c_* = 13 neurons, which is less than half the number required for the random matrix (*R_c_* ≃ 35) with the same amount of sparsity (Table S1). Note that dependence of bottleneck layer size on matrix modularity is not linear, as just 2 neurons overlapping between clusters already requires the critical bottleneck size to be at least *R_c_* = 30, making learning much harder (point #3 in Fig. 2a).

**FIG. 2.**
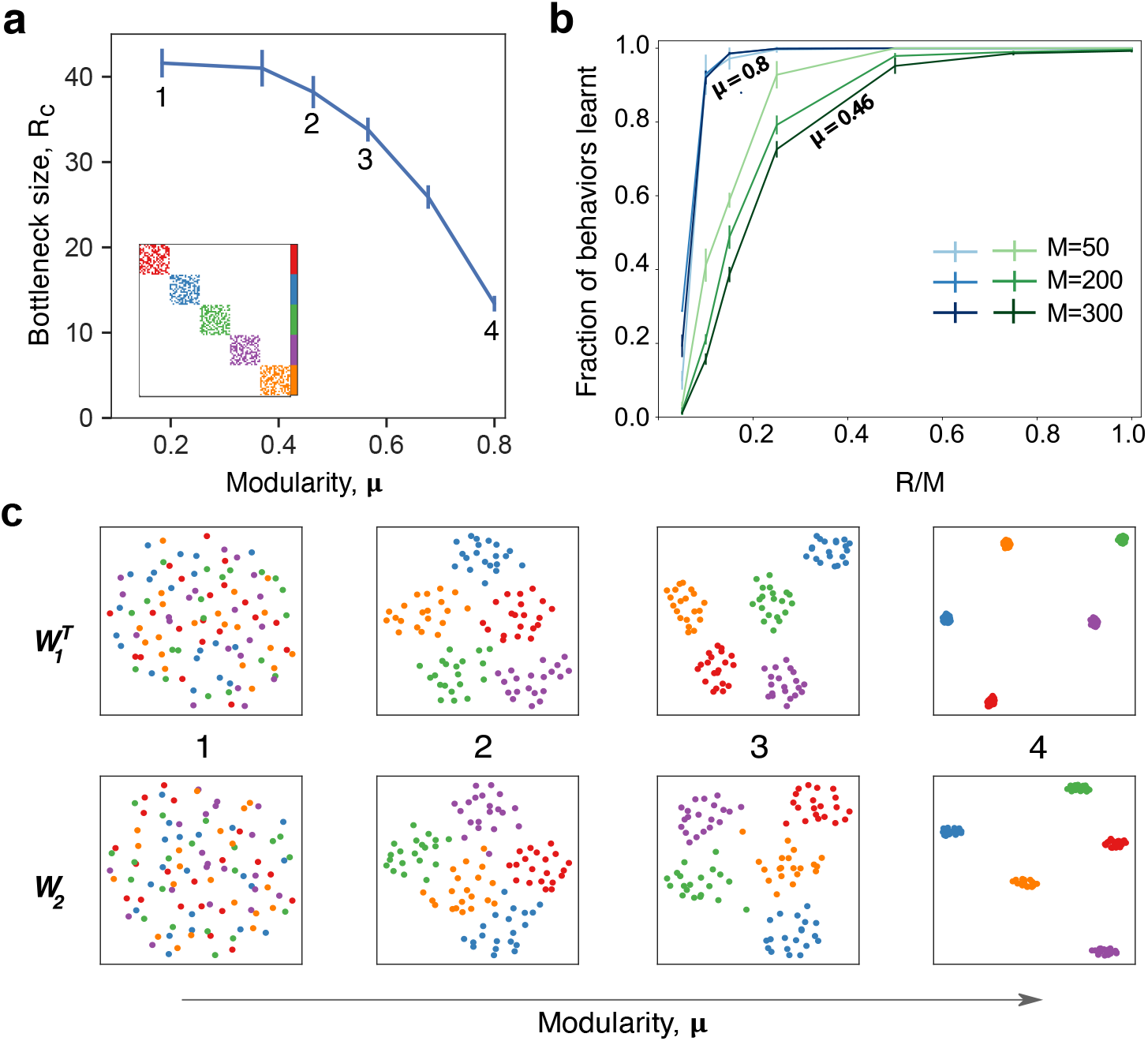
Modularity in the behavioral commands reduces critical bottleneck size and affects other network properties. **(a)** Relationship between the size of the hidden layer *R* and the modularity of the behavior matrix. Each point corresponds to a set of numerical experiments with 10 different matrices around a given modularity value (see Methods for details of data generation) for *k* = 10, *M* = *N* = 100. *R_c_* is defined as the minimal hidden layer size that was able to achieve 98% accuracy in 10^5^ epochs of training. Numbers indicate specific cases that are shown in panels b and c in more detail. Inset shows an example of behavioral command matrix for *μ* = 0.8 case (point 4). **(b)** Fraction of behaviors learned as a function of the hidden layer size, *R* for different system sizes with *N* = *M* for two levels of modularity (*μ* = 0.8 and *μ* = 0.46). Error bars correspond to the standard deviation. Results are averaged over 5 different runs with error bars corresponding to the standard deviation. **(c)** Structure of the weight matrices *W*_1_ and *W*_2_ for different modularity values. The dimensionality reduction is performed via UMAP^10^, a nonlinear method that preserves local structure in the data. The point colors correspond to the colors in the panel (a) inset: 1 (*μ* = 0.18, random matrix); 2 (*μ* = 0.46); 3 (*μ* = 0.56); 4 (*μ* = 0.8, perfectly modular matrix with 5 clusters).

In addition to making the mapping easier to learn, modularity in the behavioral matrix also helps learning scale with the system size. In Figure 2b, we plot the fraction of behaviors learned as a function of the relative size of the bottleneck layer *R* as compared to the output layer *M*, for different values of the system size (we assume *N* = *M*) and for different values of the modularity. For highly modular behavioral matrices (blue curves in Fig. 2b), we find that the size of the output doesn’t play much of a role as the bottleneck occurs when the size of the hidden layer is a similar fraction of the output sizes. On the other hand, when the behavioral commands aren’t very modular, smaller system sizes learn better for a relatively smaller bottleneck size (green curves in Fig. 2b).

Finally, we found that imposing a modular output structure also imposes a modular structure on the weights of the learned network (Fig. 2c). The modularity in the weights becomes more pronounced as the modularity of the behavioral matrix increases, similar to results found in the study of more generalized artificial neural networks^12^. Together, these results show that modularity in the behavioral matrix increases the efficiency and scaling properties of the network through creating a concomitantly modular representation within the model.

### Robustness to perturbations

Although the network is capable of reproducing behavioral commands nearly perfectly when it is near the critical bottleneck size, it might be prone to errors due to minor perturbations, including noise in the firing of the descending layer. Inspired by previous studies in flies where descending neurons were artificially activated^7,8^, we investigate the robustness of our trained neural networks by manually activating one hidden neuron at a time. We then observe the changes in the output (see Fig. 3a) to see how these activations affect the mapping between command and behavior. An example of possible outcomes on a set of behaviors under these perturbation is shown in Fig. 3c. For each behavioral command, the motor neurons can either remain unaffected - their original ‘active’ or ‘non-active’ state is maintained (grey and white pixels in Fig. 3c) or their state gets flipped – an ‘active’ neuron gets inactivated or vice-versa (red and blue pixels in Fig. 3c). The robustness of the network with respect to the activated neuron is related to the number of behaviors that are conserved, that is, behavioral commands where all activated motor neurons remain unaffected.

**FIG. 3.**
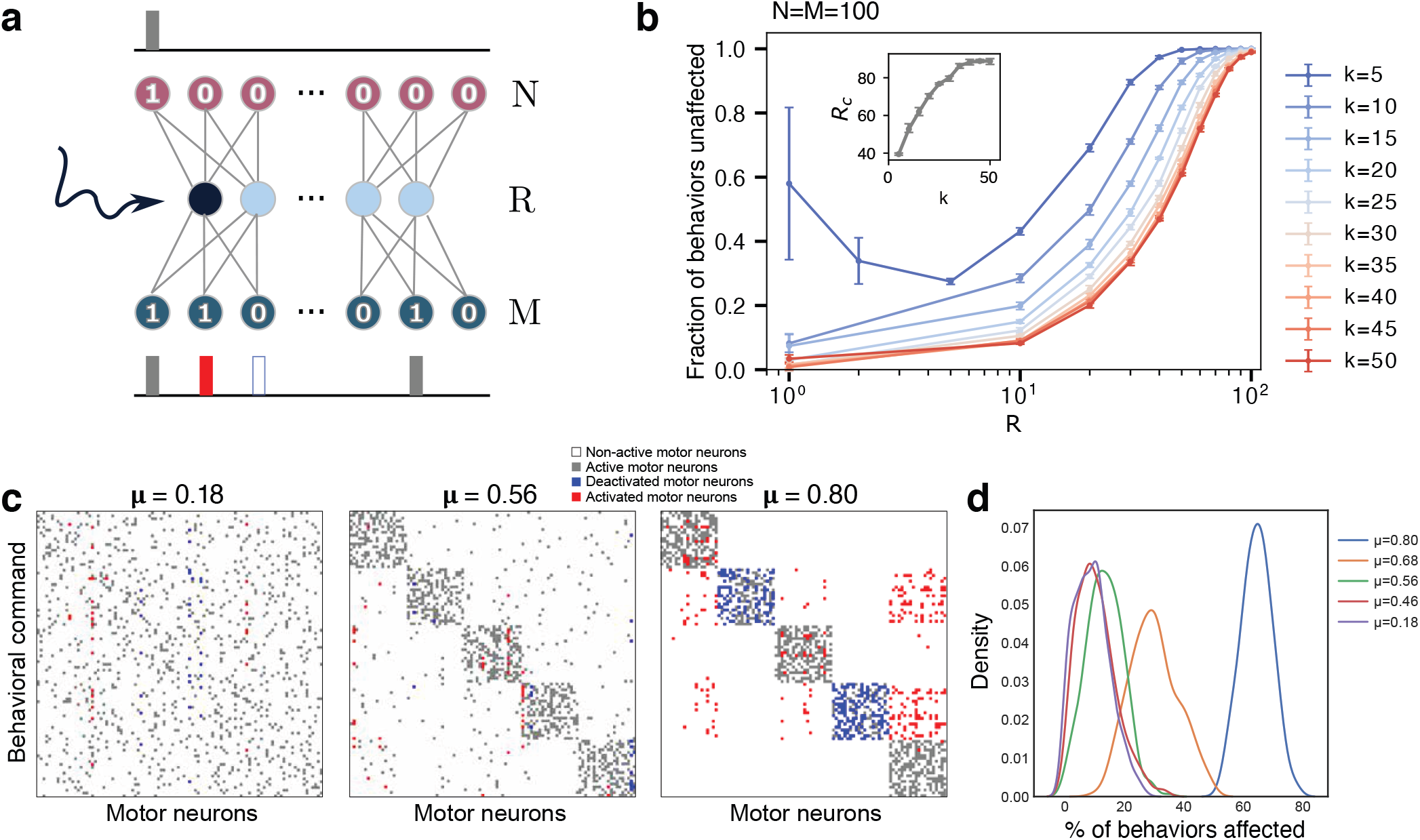
Robustness of the network to perturbations increases with the size of the hidden layer and sparsity. **(a)** Schematic of the perturbation experiment. One of the hidden neurons of the trained network is artificially forced to be on, keeping all other network parameters unchanged. The network is re-run to generate new outputs for each behavioral command. **(b)** Robustness (fraction of outputs that are unaffected by the perturbation) averaged over the effects of activating each hidden neuron as function of the hidden layer size *R*, with *N* = *M* = 100, with *k* varied from *k* = 5 to *k* = 50. The error bars are obtained by considering 10 different behavioral matrices. The inset shows the critical bottleneck size, *R_c_*, as a function of changing sparsity (varying *k*) for this network. **(c)** Example of a hidden layer perturbation on the trained networks’ behavior matrices with different modularities (all show with *R* = *R_c_*. In each case, one of the hidden neurons is kept constantly activated, while the rest of the network operates according to the trained weights. White and grey colors correspond to unperturbed motor neurons, non-active and active correspondingly. Blue indicates motor units that have been turned off, and red shows motor units that have been activated. **(d)** Distribution of the number of behavioral commands affected by the hidden layer perturbation. Colors correspond to different degrees of modularity *μ*. Each distribution was calculated based on 10 different behavioral matrices, all with *R* = *R_c_*.

Figure 3b shows the robustness of the network to these perturbations as a function of the hidden layer size *R* and varying sparsity (*N* = *M* = 100 is fixed and *k* is varied), averaged over the effects of activating each hidden neuron and each behavioral command for a randomly generated behavioral matrix (no enforced modularity). For fixed sparsity, the fraction of behaviors that are unaffected increases as the size of the hidden layer increases. At the critical bottleneck size, for example, *R_c_* = 35 for *k* = 10, 80% of behaviors were unaffected by the perturbation, indicating that the neural network has some margin of robustness. Robustness increases as we increase the intermediate layer size *R* – the behavioral commands become less sensitive to changes in each individual hidden neuron. As long as the bottleneck layer size is less than the output layer (*R < M*), networks with output signals of high sparsity (lower *k*) are more robust on average. The robustness is bounded below by the curve corresponding to maximum output signal density *k* = 50 = *M/*2. For sufficiently dense output signals 50 ≥ *k* < 5, the robustness decreases monotonically with decreasing hidden layer size for the entire range of 1 ≤ *R ≤ M*. In contrast, the robustness of high sparsity outputs (*k* = 5) decreases initially with decreasing hidden layer size, but exhibits an increase in both its mean and variance at very small hidden layer sizes (*R* < 5). This behavior is likely caused by an all-or-nothing switching relationship between the hidden neurons and the output neurons.

When applying these perturbations to more modular behavioral matrices (Fig. 3c), we find that the effects of the activations to the hidden neurons lead to more correlated changes in motor outputs. For these cases at the bottleneck size *R_c_* (which varies depending upon the modularity, see Fig. 2a), when some of the hidden neurons are activated, they not only affect a certain number of behaviors, but all of these commands tend to belong to the same cluster, which is what we would expect, given the modular structure of the weights in Fig. 2c. Moreover, activation of a neuron can lead to the complete switch from one type of behavior to the another. An example of this effect is shown in Fig. 3c. The first matrix in this panel corresponds to a random matrix of behavioral commands (also point #1 in Fig. 2a). In this case, a particular hidden neuron may be attributed to at most some set of motor neurons as its activation leads to activation of two of them and deactivation of other three. However, in the perfectly modular case, there are some neurons that are responsible for the encoding of the whole cluster (rightmost panel in Fig. 3c). When a hidden neuron is activated, it causes nearly an entire module of behaviors to be altered. This is in keeping with the previous studies showing that stimulating individual descending neurons in flies can result in dramatic behavioral effects^7,13–15^. Averaging over several behavioral matrices and perturbations (Fig. 3d), we observe that this pattern holds true in general, with more modular behavioral matrices affected more by perturbations at *R_c_*.

This effect is likely due to the different sizes of the hidden layer where the critical bottleneck size *R_c_* (the minimum number of hidden layer neurons needed to ably represent all behavioral commands) occurs, for varying levels of modularity. As the size of the hidden layer controls the susceptibility towards perturbations (Fig. 3b), highly modular behavioral matrices that have a much smaller *R_c_* (Fig. 2a), are affected to a larger extent by the perturbations. In general however, if the size of the hidden layer is fixed, modularity *increases* the robustness to perturbations (Fig. 4a, Fig. S4a).

**FIG. 4.**
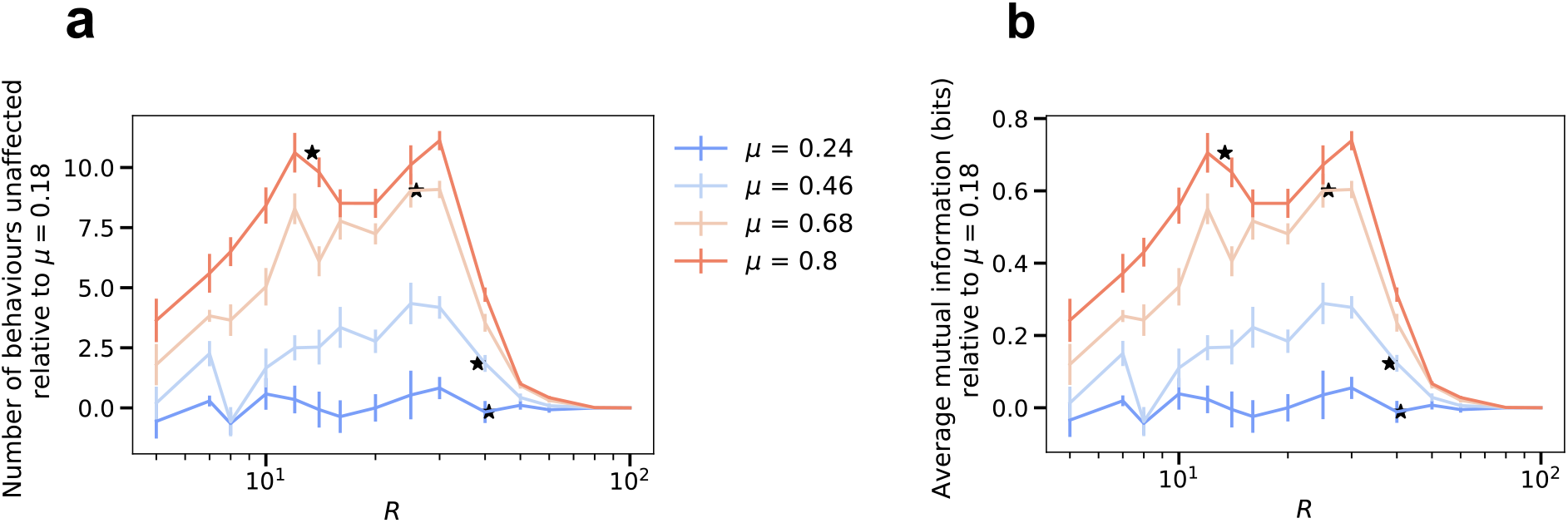
Modularity improves robustness to perturbation and increases context dependency for a fixed size of the hidden layer. To highlight the effects of increasing modularity, we show the results relative to the lowest modularity, *μ* = 0.18 as 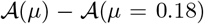, where 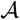 is the robustness of the network (panel **a**) and the mutual information (panel **b**) defined as follows. The robustness 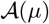 is defined as the numbers of behaviors that are not affected upon forcefully activating a neuron in the network. The figure for the absolute values is reported in the Supplementary Material, Fig. S4. The value of *R_c_* for each modularity value is shown as stars. **(a)** Robustness of the network averaged over the effects of activating each hidden neuron as a function of the hidden layer size, R and varying levels of modularity, *μ*. **(b)** Average mutual information (defined as in Eq. 7) between the input and output distributions after forced activation of each hidden neuron as a function of the size of the hidden layer R, and varying levels of modularity, *μ*. The mutual information turns out to be 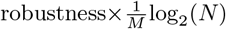 due to the absence of stereotypy. **(a)-(b)** *N* = *M* = 100 and results are means over 5 iterations with the error bars corresponding to the standard deviation.

Thus, when constrained by a fixed size of the hidden layer, increasing the modularity and sparsity of the behavioral commands helps increase the robustness of the network to artificial perturbations. However, robustness suffers if the goal is to operate the network at the smallest possible critical bottleneck size for a given number of behavioral commands.

### Context dependency of behaviors

Previous experimental studies in fruit flies observed that optogenetically activated behaviors in flies often depend on their behavioral state prior to activation^7,8^. This effect can be quantified by calculating the mutual information between the distribution of a fly’s behaviors before and after artificial neural activation. We refer to this effect as *context dependency*. In order to understand this experimentally observed effect within the framework of our model, we calculated the mutual information between the input and output distributions in the presence of an activated hidden neuron, while varying the size of the hidden layer and modularity (Figs.4b and S4b, see Methods for details). This calculation provides a measure of how much information about the input distribution is contained in the output distribution in the presence of artificial activation.

With the input distribution corresponding to the fly’s intended behavioral output (the one-hot encoded initial layer from Fig.1a) and the modified output corresponding to the set of behaviors that the artificial activation triggers, we see that increasing the bottleneck constraint (reducing *R*) lowers the overall mutual information - thus, it becomes harder to predict what the triggered behavior will be. On the other hand, a higher amount of modular structure in the output behavioral commands increases the mutual information for a fixed size of the hidden layer, with a maximum increase of around 0.8 corresponding to about a 30% increase between the two extreme values of modularity (*μ* = 0.18 and *μ* = 0.8) considered here. Thus, our model predicts that increased modular structure in the behavioral matrix not only increased robustness to perturbations (for a given *N*, *M*, and *R*), but also results in increased context dependency. These results are consistent with the finding of context dependency and modularity in the *Drosophila* VNC.

It is worth mentioning that we find that the mutual information is proportional to the robustness (Figs.4a and 4b) with a proportionality constant 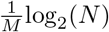 (see Methods). This is a consequence of an absence of stereotypy in our simplified model, multiple inputs don’t get mapped to the same output on forced activation.

Given these results, we explored what predictions our model makes for two additional types of perturbation experiments that have not, to our knowledge, been systematically performed. First, we asked what the effects would be for deactivating, rather than activating, individual hidden layer neurons (Fig.5a). As one might expect for a binary encoded network, the effect of deactivating individual neurons on the robustness of the network is qualitatively similar to that for activation. The network is more robust to the perturbation as the size of the hidden layer increases. For any given size of the hidden layer, modularity increases the network’s robustness to deactivating perturbations.

**FIG. 5.**
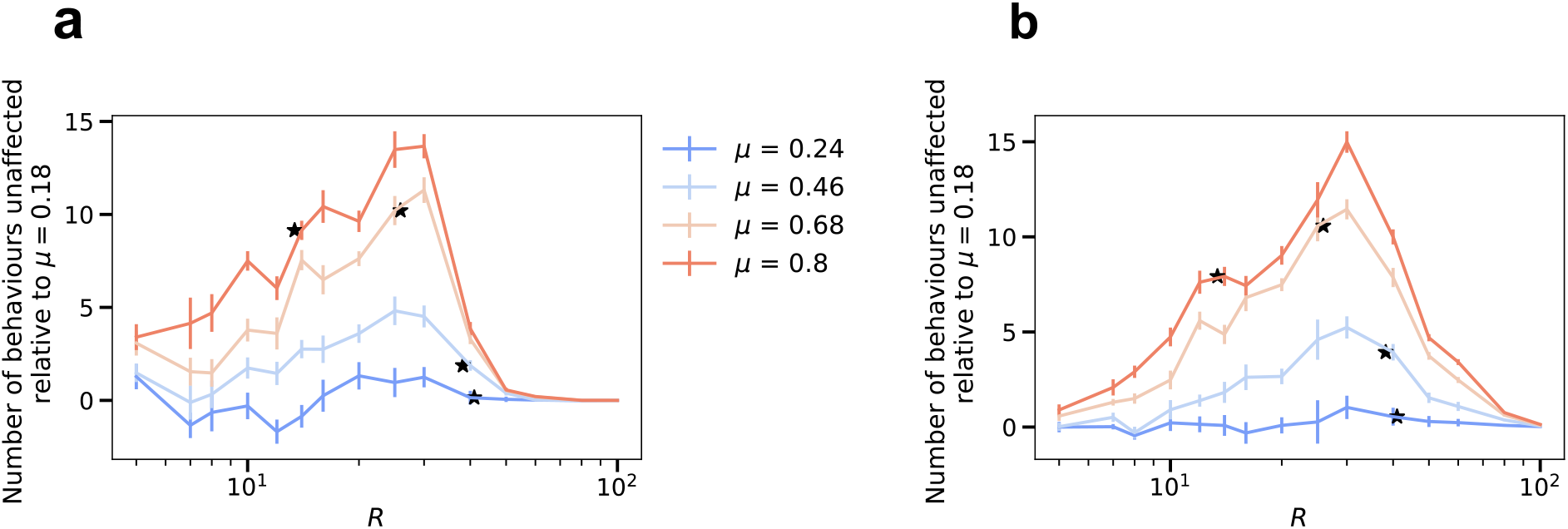
Future excitation and inhibition experiments predict modularity is always associated with improved robustness. To highlight the effects of increasing modularity, we show the results relative to the lowest modularity, *μ* = 0.18 as 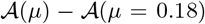, where 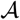 is the robustness of the network upon de-activating each hidden neuron (panel **a**) and the robustness upon activating pairs of hidden neurons one at a time (panel **b**) defined as follows. The figure for the absolute values is reported in the Supplementary Material, Fig. S5. The value of *R_c_* for each modularity value is shown as stars. **(a)** Robustness of the network averaged over the effects of de-activating each hidden neuron as a function of the hidden layer size, R and varying levels of modularity, *μ*. **(b)** Robustness of the network averaged over the effects of activating a pair of hidden neurons as a function of the hidden layer size, R and varying levels of modularity, *μ*. **(a)-(b)** *N* = *M* = 100 and results are means over 5 iterations with the error bars corresponding to the standard deviation.

Similarly, we also explored whether activating pairs of hidden layer neurons (rather than individual neurons) leads to increased context dependency with modularity as well (Fig.5b). We find similar results in this case (averaging over all possible pairs of hidden layer units across many networks).

## DISCUSSION

Understanding how animals use their nervous system to control behavior is one of the key questions in neuroscience. A key component of most animal’s nervous system is an information bottleneck between cognitive decision-making in the brain and the neurons that are responsible for the performance of behaviors. In this work, we use a simple feed-forward neural network to understand the consequences of having such a bottleneck and identify different aspects of the network architecture that can still enable robust learning despite having such a constraint. For each set of network parameters, we identify the smallest size of the hidden layer (bottleneck size) that still allows near perfect learning. We find that increasing the sparsity of the output behavioral commands reduces this bottleneck size and increases the robustness of the network.

In addition to sparsity, we find that an increased modularity in the behavioral commands helps to reduce the bottleneck size and increases robustness. This observation could provide an explanation for why such a modular structure has evolved in the behavioral commands in animals, so far observed in flies. Our simple model is also able to predict the experimentally observed context dependency between behavioral states before and after the forced activation of hidden neurons. We find that lowering the size of the hidden layer reduces context dependency, but context dependency increases with increasing modularity for a fixed hidden layer size. Overall, the modular nature of the output makes it easier for the network to learn in the presence of a bottleneck, increases it’s robustness but also leads to a higher amount of context dependency.

This model described here is obviously simplistic in architecture and dynamics (in that it lacks them) and is highly unlikely to accurately describe the dynamical activity of ventral nerve cord function, where recurrent connections and temporal structure are important features of the system’s functioning^6,16^. Future work would incorporate the effects of temporal dynamics, as well as using more biophysically realistic neurons. However, despite its simplicity, our model recapitulates several nontrivial features that are observed in experiment, and makes predictions as to the effects of artificially inhibiting neurons or of simultaneously stimulating multiple neurons, allowing for general principles of information-limited motor control to be elucidated and new hypotheses to be tested.

## METHODS

### Network architecture and training

To mimic the structure of the neural chord, we built a feed-forward fully-connected neural network with one hidden layer (see Fig. 1). The network is constructed with the Python framework PyTorch. The input layer represents decision neurons of number *N* : they send the signal from the brain down the network leading to a certain behavioral output. The hidden layer of size *R* represents descending neurons of the neural chord: it transmits the signal down to the motor neurons, which are the output layer of the network of size *M*. We used the sigmoid as our activation function, serving as an approximation of the transmission of the neural signal. The functioning of the neural network can be understood explicitly from its mathematical definition. The first layer applies a linear transformation on the input sequence 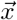 via the weight matrix, 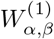 connecting neuron *α* in the first layer with neuron *β* in the following equation,

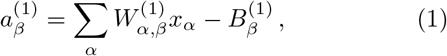

while the second and last layer applies the activation function *ρ*(**a**) on **a**^(1)^ as,

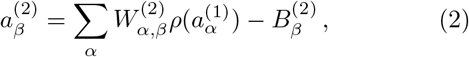

with *ρ*(**a**) given by the sigmoid *ρ*(*x*) = 1/(1 + *e*^−*x*^) and **B**^(1)^ (**B**^(2)^) is the bias, an additive constant. The output of the network is defined as *f* (**x**, **W**) **a**^(2)^, where **W** contains all the parameters, comprising the biases.

We fixed the size of the input layer (*N* = 100) throughout our experiments, while varying the sizes *R*, *M* of the hidden and output layers. We trained the network in the following fashion: we fixed the input and output matrices, i.e. decision and behavior matrices respectively; we trained the network in a feed-forward manner using stochastic gradient descent with momentum and used the mean-squared error (MSE) loss function to assess learning performance; we stopped training after 10^5^ epochs, which corresponds to when the loss curve flattens and the network is no longer learning. The output **y**= *f* (**x**, **W**) of the trained network is then binarized by rounding each entry and the trained weights and biases defining the network are saved for further analysis. Along with these parameters, the number of behaviors learnt, obtained by comparing each entry of the output **y** with the imposed behavior, is also stored.

### Modularity

We use the NetworkX 2.5 Python package to calculate modularity using the function ‘networkx.algorithms.community.modularity’ by treating the output matrix of behavioral commands as an adjacency matrix of a graph. Here modularity is defined as^11^,

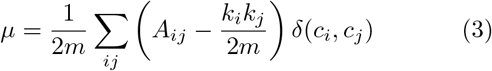

where *m* is the number of edges, *A_ij_* is the adjacency matrix, *k_i_* is the degree of *i* and *δ*(*c_i_*, *c_j_*) is 1 if *i* and *j* are in the same community and 0 otherwise.

### Data generation

The input data for all of our numerical experiments is always a 100 × 100 identity matrix. Each row of this matrix corresponds to the signal of performing one behavior from the output matrix. We generated several sets of output behavior matrices. In Fig. 1, we varied the sparsity of the output matrix by changing the number of randomly activated units in a given row, i.e. the number of 1s. In Fig. 3 we generated modular behavior matrices by introducing dense and sparse clusters into the output matrix. We start with 5 perfect clusters, i.e. no activated units are in common between 2 different clusters. Then, we generate matrices with different degree of modularity by deactivating some of the units within the cluster and activating the same number of units outside of the cluster so that the sparsity is preserved. In each case we generated 10 different behavior matrices for statistical purposes.

### Checking the robustness of the network

We checked the robustness of the network by forcefully activating one of the hidden layer neurons. This is achieved by setting its corresponding weight in the first weight matrix W^(1)^ to an arbitrarily high value. We propagate the input matrix through the resulting perturbed network to get an output behavior matrix to be compared to the original output. In this way we can monitor how many of the original output behaviors were changed by the forceful activation. These steps are repeated for each individual hidden neuron and the results are averaged over the number of hidden neurons.

### Mutual information calculation

Mutual information (MI) between two distributions is the measure of the amount of information one distribution has about the other. For two discrete binary random variables *X* and *Y* embedded in ℝ^*N*^ with joint distribution *P* (*X, Y*) it is given by^17^,

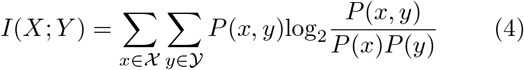

where *P* (*X*) and *P* (*Y*) are the marginal distributions. In the absence of forced activation, the perfect learning case has a one to one mapping between the input and output distributions and hence the MI is log_2_*N*. This perfect mapping gets perturbed on forced activation which can lead to one of the three different scenarios: (i) the input-output mapping is still unaffected, (ii) the input gets mapped to another output (stereotypy), and (iii) the input gets mapped to a completely different output that is not part of the original output distribution. This last case suggests that the input possess no information about the output.

Suppose we have *N* inputs *x* and *M* outputs *y* where we assume that they follow a uniform distribution, that is, *P*(*x*) = 1/*N* and *P* (*y*) = 1*/M*. After forced activation, let *n_i_* be the number of inputs associated with each output *y_i_* where *n_i_* ≥ 0. This gives us 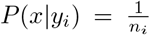 when *n_i_* > 0 and *P* (*x*×*y_i_*) = 0 when *n_i_* = 0. The mutual information then reads

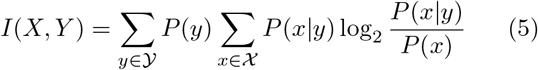

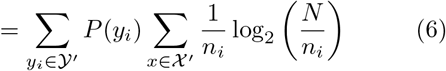

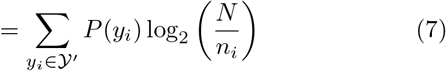

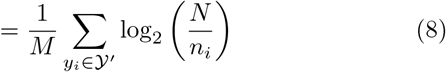

where 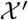 is the set of *n_i_* inputs associated with each output 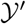, Y is the set of *m* outputs with *n_i_* > 0.

### Statistical analysis

Error bars in the figures are standard deviations that were calculated by averaging simulation results for 10 different output matrices unless specified otherwise.

We used the UMAP^10^ method to visualize the structure in weight matrices.

## Code availability

The code for both our simulations and statistical analysis, can be downloaded from: https://github.com/drahcir7/bottleneck-behaviors.

## Acknowledgements

The authors thank the organizers of the 2019 Boulder Summer School for Condensed Matter and Materials Physics for the opportunity to meet and start a collaboration on this project. G.J.B. was supported by the Simons Foundation and a Cottrell Scholar Award, a program of the Research Corporation for Science Advancement (25999). A.N. was supported by a grant from the US National Institutes of Health (DP5OD019851). G.H.Z. acknowledges support from the Paul and Daisy Soros Fellowship and the National Science Foundation Graduate Research Fellowship under Grant No. DGE1745303. R.R. was supported by the Swiss National Science Foundation under grant No. 200021-165509/1.

## Author contributions

G.J.B. initiated the project and supervised the study; A.N., G.Z., V.D. and R.R. performed simulations and analysis. All authors devised the study and wrote the paper.

## Supplementary Figures

**FIG. S1.**
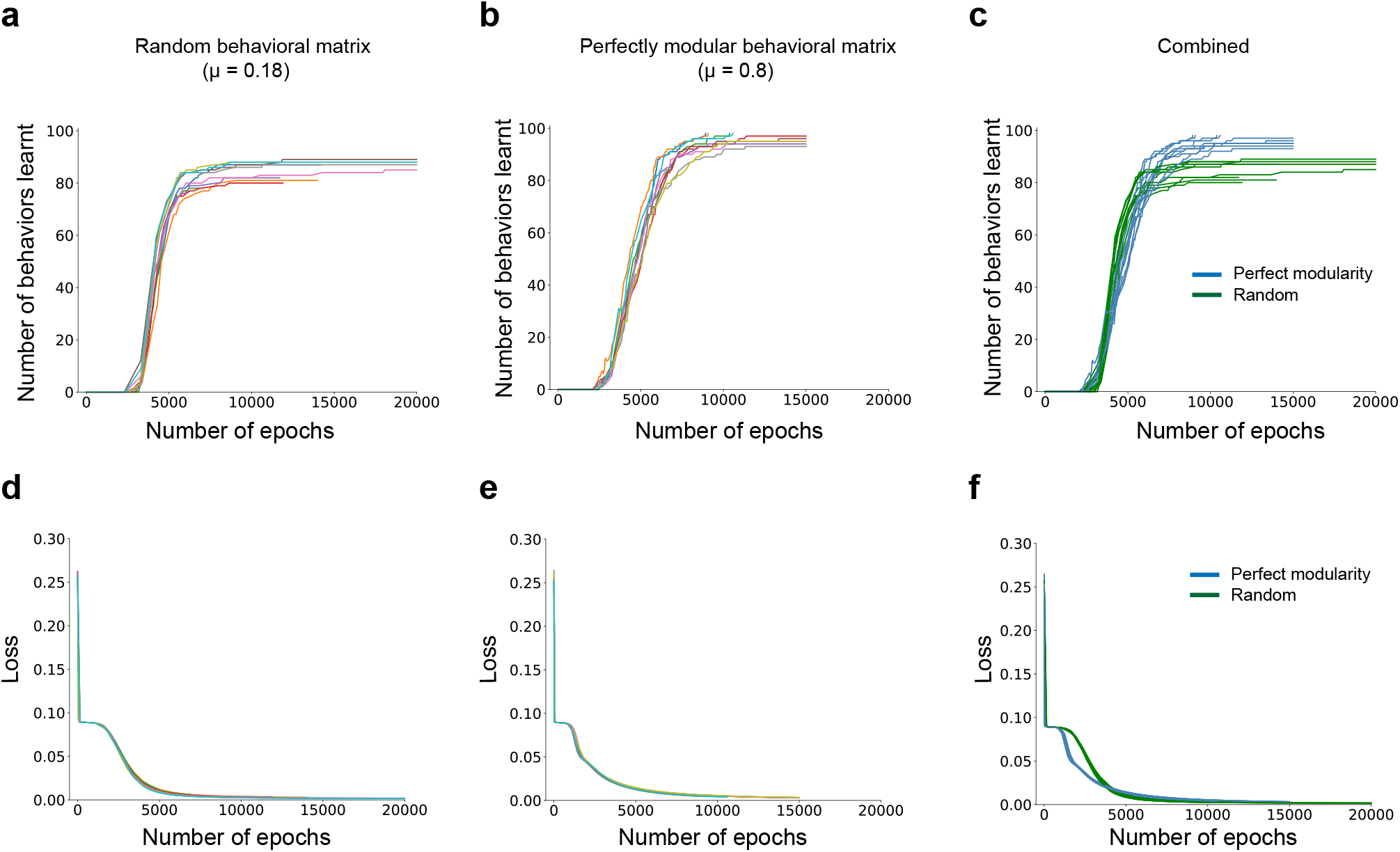
Learning **(a)-(c)** and loss curves **(d)-(f)** obtained while training the network when the behavioral matrix is **(a)&(d)** random (*μ* = 0.18), **(b)&(e)** perfectly modular (*μ* = 0.8) with the combined results in **(c)&(f)**. *N* = *M* = 100 with *R* = 25 for the random and *R* = 13 for the fully modular case. For each value of modularity we plot n=10 curves.

**FIG. S2.**
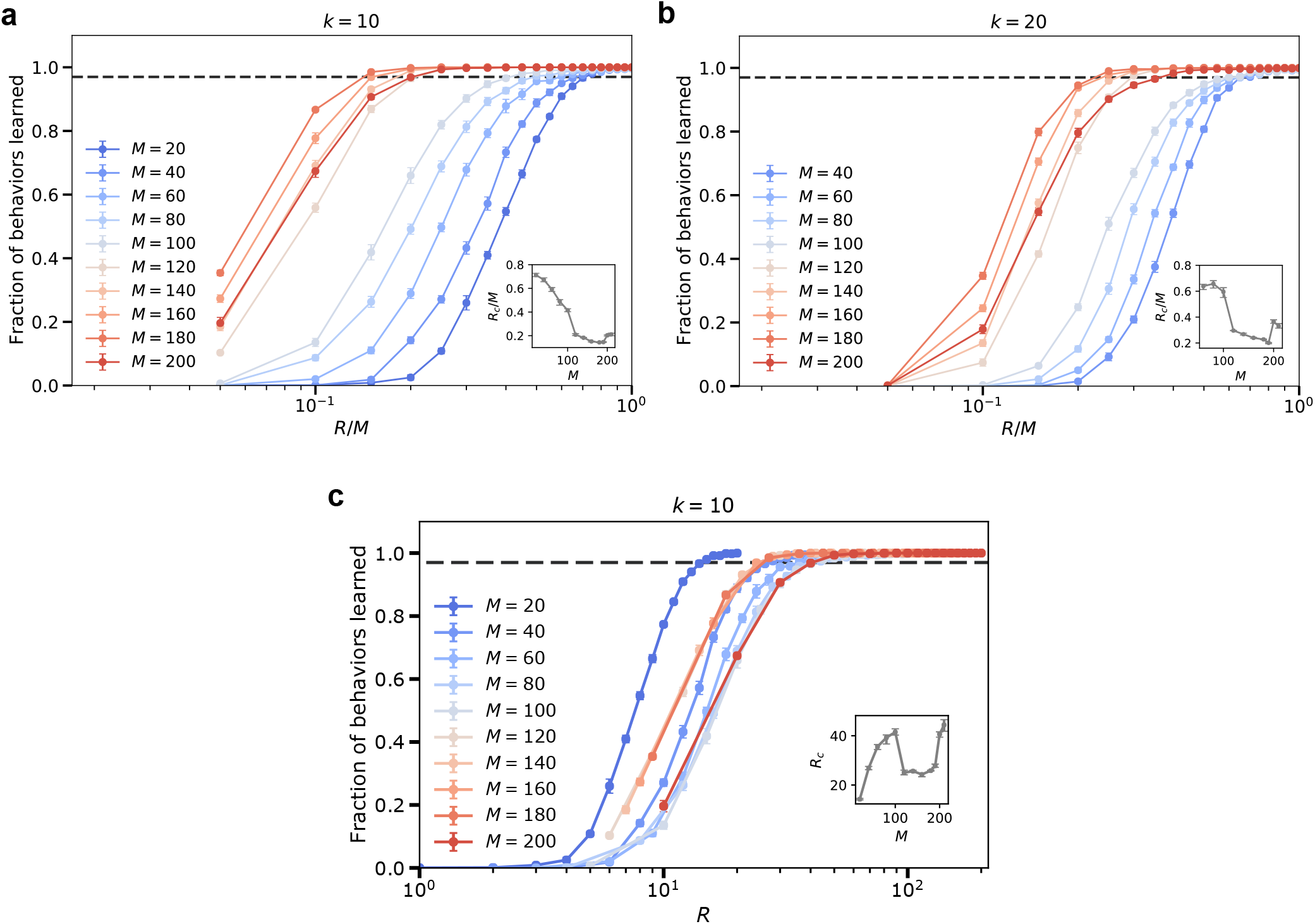
**(a)-(b)** Fraction of behaviors learned as a function of hidden layer size *R* and fixed input layer size *N* = 100 for varying output layer size *M* and fixed number of active output neurons (*k* = 10 (a) and *k* = 20 (b)). The horizontal axis is the hidden layer size scaled by the output layer size *R/M*. Insets show the scaled critical bottleneck size *Rc/M* as a function of *M*. Each point is averaged over 30 random input-output combinations. Dashed line indicates critical bottleneck threshold. **(c)** Subfigure (a) with the horizontal axis replaced by the unscaled hidden layer size *R*. The curves loosely collapse onto each other; this allows us to use one value of *M* for most of this work. The inset shows the critical bottleneck size *R_c_* as a function of *M*, with *M* = 100 as the output layer size for which the network has most difficulty learning (highest threshold bottleneck size *R_c_*). We thus focus our studies on *M* = 100 networks.

**FIG. S3.**
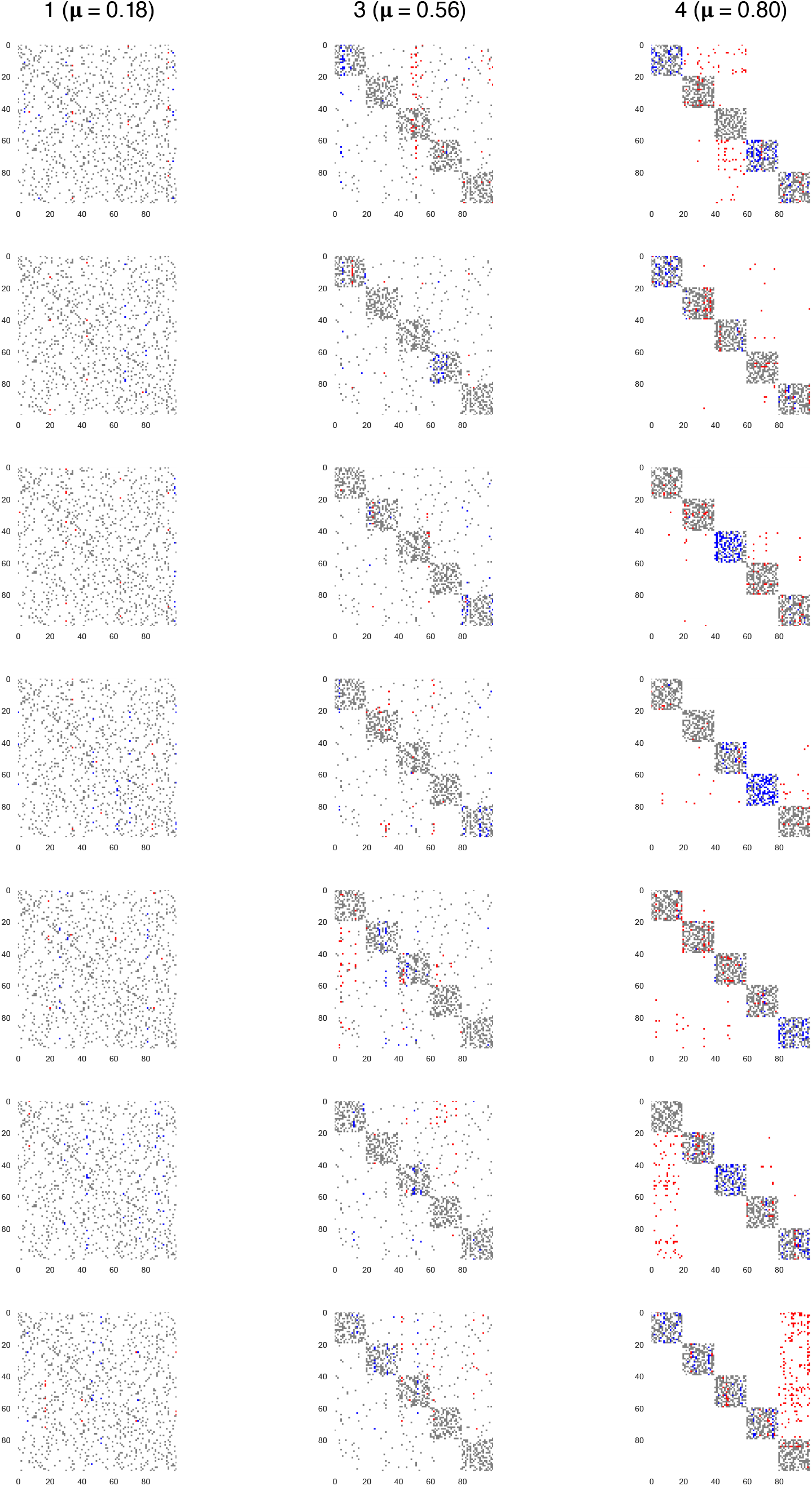
Other examples of a hidden layer perturbation on the networks trained for behavior matrices with different modularities at *R_c_* point. Numbers correspond to the point numbers in panel a. In each case one of the hidden neurons is kept constantly activated, while the rest of the network tried to reproduce the original set of behaviors. White and grey colors correspond to unperturbed motor neurons, non-active and active correspondingly. Blue indicates motor units that have been deactivated for different behaviors due to this hidden neuron activation. Conversely, red shows motor units that have been activated.

**FIG. S4.**
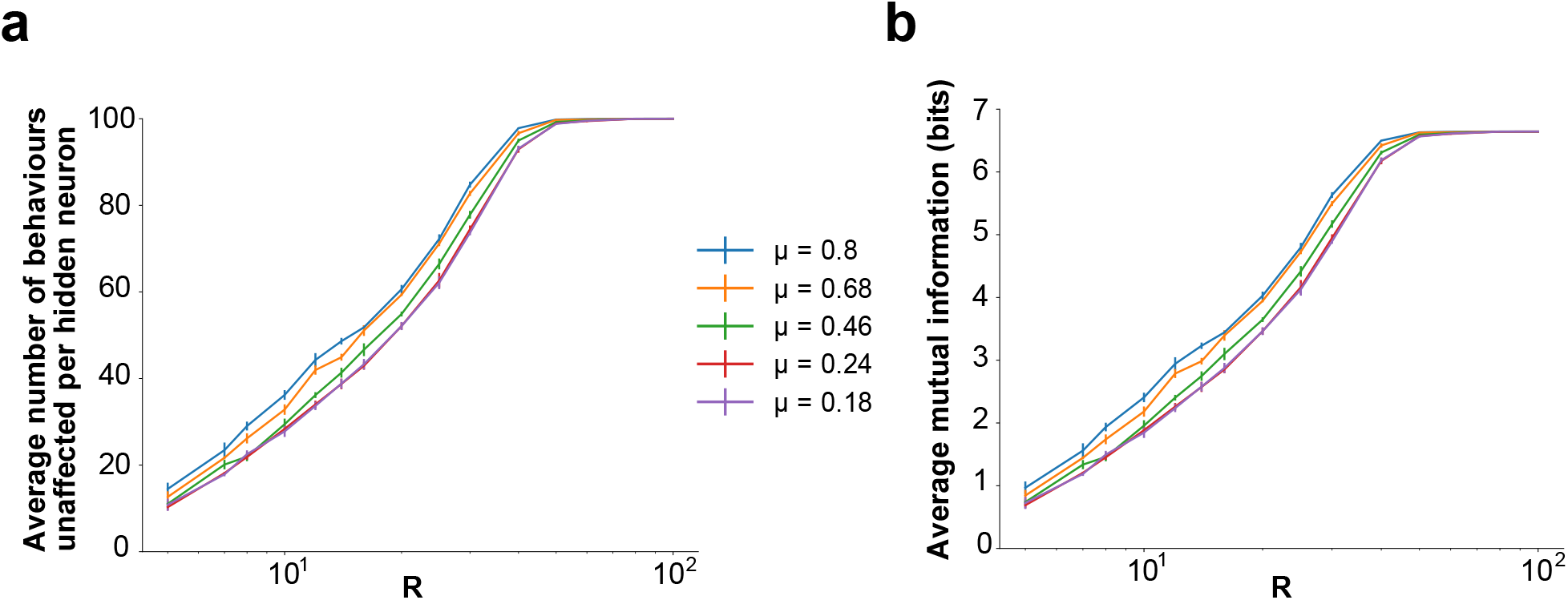
**(a)** Robustness of the network averaged over the effects of activating each hidden neuron as a function of the hidden layer size, R and varying levels of modularity, *μ*. **(b)** Average mutual information between the input and output distributions after forced activation of each hidden neuron as a function of the size of the hidden layer R, and varying levels of modularity, *μ*. **(a)-(b)** *N* = *M* = 100 and results are means over 5 iterations with the error bars corresponding to the standard deviation.

**FIG. S5.**
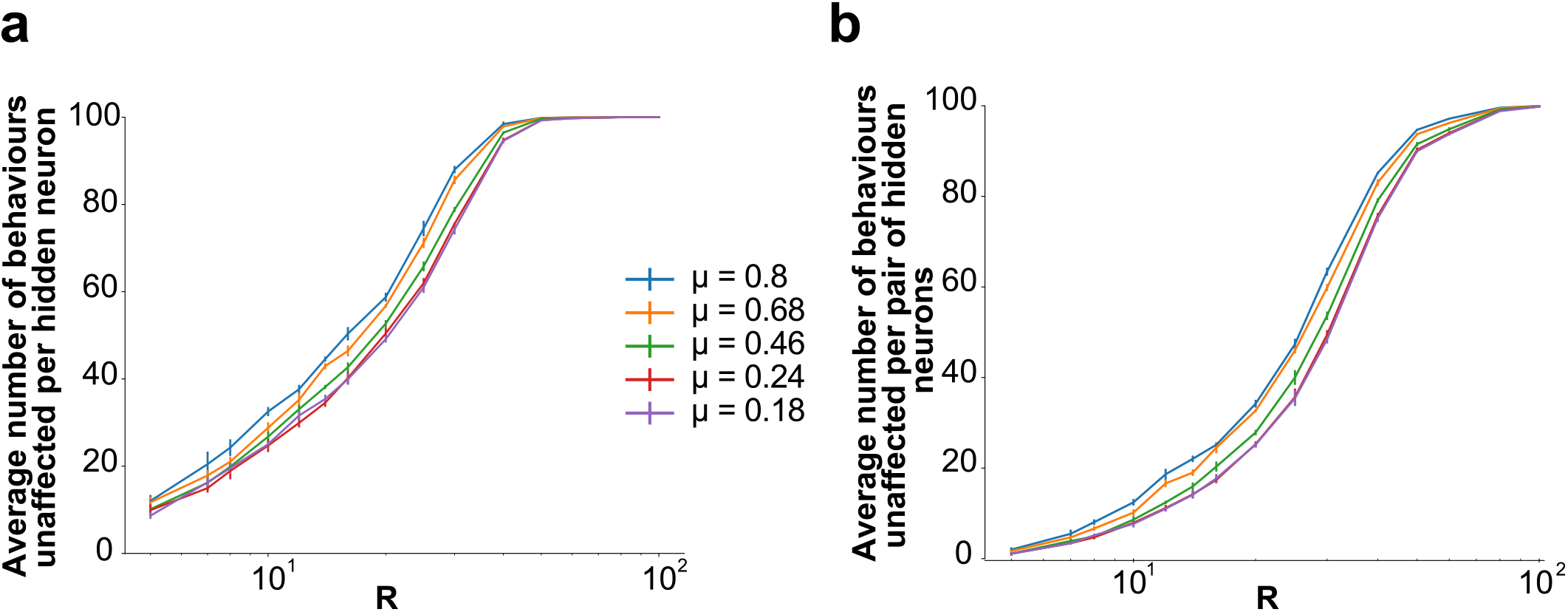
**(a)** Robustness of the network averaged over the effects of de-activating each hidden neuron as a function of the hidden layer size, R and varying levels of modularity, *μ*. **(b)** Robustness of the network averaged over the effects of activating a pair of hidden neurons as a function of the hidden layer size, R and varying levels of modularity, *μ*. **(a)-(b)** *N* = *M* = 100 and results are means over 5 iterations with the error bars corresponding to the standard deviation.

## Supplementary Tables

**TABLE S1.** Values of the critical bottleneck size *R_c_* for different values of modularity *μ*. The mean and standard deviation are calculated over 10 experiments with different output matrices for each value of modularity.

| $\mu$ (mean ± std) | $R_c$ (mean ± std) |
| --- | --- |
| $0.180 \pm 0.004$ | $41.6 \pm 1.7$ |
| $0.369 \pm 0.003$ | $41.0 \pm 2.1$ |
| $0.464 \pm 0.003$ | $38.2 \pm 1.9$ |
| $0.566 \pm 0.002$ | $33.8 \pm 1.4$ |
| $0.677 \pm 0.002$ | $25.9 \pm 1.4$ |
| $0.800 \pm 0.0001$ | $13.4 \pm 0.9$ |

